# Antiviral and anti-inflammatory effects of Tabamide A derivative, TA25, against human rhinovirus and multiple zoonotic viruses *in vitro* and *in silico*

**DOI:** 10.1101/2025.11.23.690071

**Authors:** Bohyeon Kim, Altanzul Bat Ulzii, Taehun Lim, Jin Woo Kim, Wonkyun Ronny Im, Shivani Rajoriya, Augustine Natasha, Divya Misra, Hennisa Hennisa, Na Young Kwag, Daewoon Yoon, Gyunghee Jo, Young Bin Park, Jinkyu Park, Jeong Tae Lee, Sung Ho Jeon, Won-Keun Kim

## Abstract

Human rhinovirus (HRV), first isolated in 1956, belongs to the family *Piconaviridae* containing a positive-sense, single-stranded RNA genome. HRV causes mild cold and severe respiratory disease, such as asthma, COPD, and pneumonia. To date, no Food and Drug Administration-approved antiviral or anti-inflammatory drugs are available for HRV. TA25 is a phenolic amide derivative extracted from the leaves of *Nicotiana tabacum*. To investigate the potential candidate for antiviral therapeutics against zoonotic viruses, we evaluated the antiviral potency of TA25 for HRV and multiple zoonotic viruses. The antiviral and anti-inflammatory effects of TA25 were evaluated using RT-qPCR and RNA-seq. Strand-specific RT-qPCR was performed to measure genomic and anti-genomic RNA expression after TA25 treatment. In addition, an AI-based docking test was conducted to investigate the binding affinity of TA25 with viral target proteins. TA25 induced a significant reduction in viral replication and suppressed the expression of pro-inflammatory genes. Inhibition of viral replication by TA25 treatment was confirmed by strand-specific RT-qPCR. TA25 showed broad-spectrum antiviral activity against multiple viruses, including HRV-1A, Zika virus, Dengue virus, Vaccinia virus, and Influenza B virus Victoria. Using an AI-driven structure-based docking analysis, TA25 showed the strongest binding affinity with the HRV 2B protein. This study demonstrates that TA25 confers the broad antiviral and anti-inflammatory activity against HRV and multiple zoonotic viruses. These findings provide valuable insights into antiviral strategies of TA25 for a promising therapeutic candidate in response to emerging RNA and DNA viruses.

## 1. Introduction

Broad-spectrum antiviral agents are a crucial strategy for preventing and rapidly intervening the outbreaks against emerging and re-emerging viruses (1). In recent years, multiple outbreaks of emerging infectious diseases have occurred as a consequence of zoonotic spillover of viruses from animal reservoirs to humans, including Zika virus (ZIKV), avian influenza, severe acute respiratory syndrome coronavirus 2 (SARS-CoV-2) and Monkeypox virus (MPXV) infection (2). The continuous threats posed by zoonotic viruses emphasize the urgent need for the development of effective antiviral therapeutics. The advancement of broad-spectrum antiviral therapeutics remains a critical and sustainable focus in academic research and pharmaceutical development. Infections of Flaviviruses, such as ZIKV and dengue virus (DENV) have shown increasing incidence and persistence as global public health challenges (3). ZIKV was introduced in Brazil where it caused more than one million cases in 2015, followed by spreading to 84 countries (4). DENV infections were reported, with an approximately 390 million cases of infections worldwide. Of these, 96 million showed mild-to-severe symptoms with annual 25,000 deaths (5). Vaccinia virus (VACV) belongs to same family with MPXV that has emerged as one of the most significant infectious diseases caused by a poxvirus (6). The unexpected non-endemic outbreak in 2022 is considered as a new global threat after coronavirus disease 2019 (COVID-19) pandemic. The fatality rate of MPXV infection ranges from 1% to 10%. Despite of their clinical and epidemiological significance, there are no Food and Drug Administration (FDA)-approved antiviral therapeutics against these zoonotic viruses. Compounds isolated from *Nicotiana tabacum*, a stout herbaceous plant cultivated worldwide and a commercial source of tobacco have demonstrated potent antiviral activities, including inhibition of human immunodeficiency virus-1 (HIV-1), tobacco mosaic virus (TMV), and cytotoxicity activities (7). Tabamide-A (TA) is a phenolic amide isolated from the leaves of *Nicotiana tabacum* and exhibited antiviral activity. TA showed high anti-TMV activity, with an inhibition rate of 38.6%, which was higher than that of the positive control, ningnanmycin (8). Moreover, TA showed antiviral activity against influenza A viruses, with an EC_50_ of 2.72 μM in both *in vitro* and *in vivo* models (9). These findings underscore the broad-spectrum antiviral potency of TA against clinically significant respiratory viruses.

Human rhinovirus (HRV), a common cause of common cold, was first isolated by Dr. Winston Price from Johns Hopkins University in 1956 (10). HRV is a positive-sense, single-stranded, non-enveloped RNA virus (11). In host cells, viral RNA is translated using the host’s ribosomal machinery facilitated by viral protein VPg (12). After translation, VPg undergoes uridylation and is translocated to the 3’-end of the genomic RNA, specifically to the poly-A tail region, to initiate viral replication. VPg serves as a primer for synthesis of a negative-sense of anti-genomic RNA (13). The newly synthesized anti-genomic RNA functions as a template for the generation of positive-sense RNA that functions as both genomic RNA and mRNA (14).

HRV is a one of the respiratory viruses that causes the severe airways diseases such as asthma and COPD (15). However, no FDA-approved or commercialized anti-HRV drugs are currently available (16). Several antiviral drug candidates have been investigated such as rupintrivir, an HRV 3C protease inhibitor. *In vitro* studies have demonstrated the efficacy of rupintrivir, and *in vivo* studies have confirmed that rupintrivir has a good safety profile and moderate reductions in common cold symptoms and viral titres. However, owing to its limited clinical benefits, further trials on rupintrivir have not been performed (17). These findings highlight the urgent need for novel and effective antiviral agents against HRV.

In this study, we evaluated the antiviral activity of TA25 against HRV and multiple viruses *in vitro*. The reduction in the expression of immune and inflammatory genes induced by HRV were confirmed in TA25-treated cells. Broad-spectrum antiviral activity was also confirmed against RNA viruses such as ZIKV, DENV, Influenza B virus Victoria lineage (IBV Victoria), and a DNA vaccinia virus (VACV). Furthermore, we employed AI-driven structure-based docking analysis to elucidate the possible mechanism of TA25 for inhibiting viral replication. Therefore, our results highlight TA25 as a broad-spectrum antiviral therapeutic for developing effective antiviral against HRV and zoonotic viruses.

## 2. Results

### 2.1. TA25 shows the antiviral activity of HRV-B14 in various treatment conditions and dose-dependent manner

To evaluate the antiviral activity of TA25 against HRV-B14, viral RNA expression levels and plaque formation were analyzed by the treatment after viral infection (post-treatment), before infection (pre-treatment), and simultaneously with infection (co-treatment). In the post-treatment condition, TA25 significantly reduced viral replication in a dose-dependent manner, with approximately a 100-fold reduction at a 30 μM concentration (Figure 1A). A corresponding 100-fold decrease in infectious viral particles was also observed (Figure 1B). The EC_50_ of TA25 in the post-treatment condition was 3.88 μM (Figure 1C). In the pre-treatment condition, a dose-dependent antiviral activity of TA25 was also shown, with approximately a 100-fold reduction in viral RNA and particles at 30 μM (Figures 1D and 1E). The EC_50_ was 4.97 μM (Figure 1F). In the co-treatment condition, similar dose-dependent antiviral activity was observed, with a 100-fold reduction at 30 μM (Figures 1G, 1H). The EC_50_ under co-treatment was 11.69 μM (Figure 1I). The half-maximal cytotoxic concentration (CC_50_) of TA25 was 270.2 μM (Figure S1A). The selectivity index (SI), the ratio of the toxic concentration of TA25 to its antiviral efficacy, was 69.64 in the post-treatment. SI was 54.37 and 23.11 in the pre- and co-treatment, respectively (Figure S1B).

**Figure 1.**
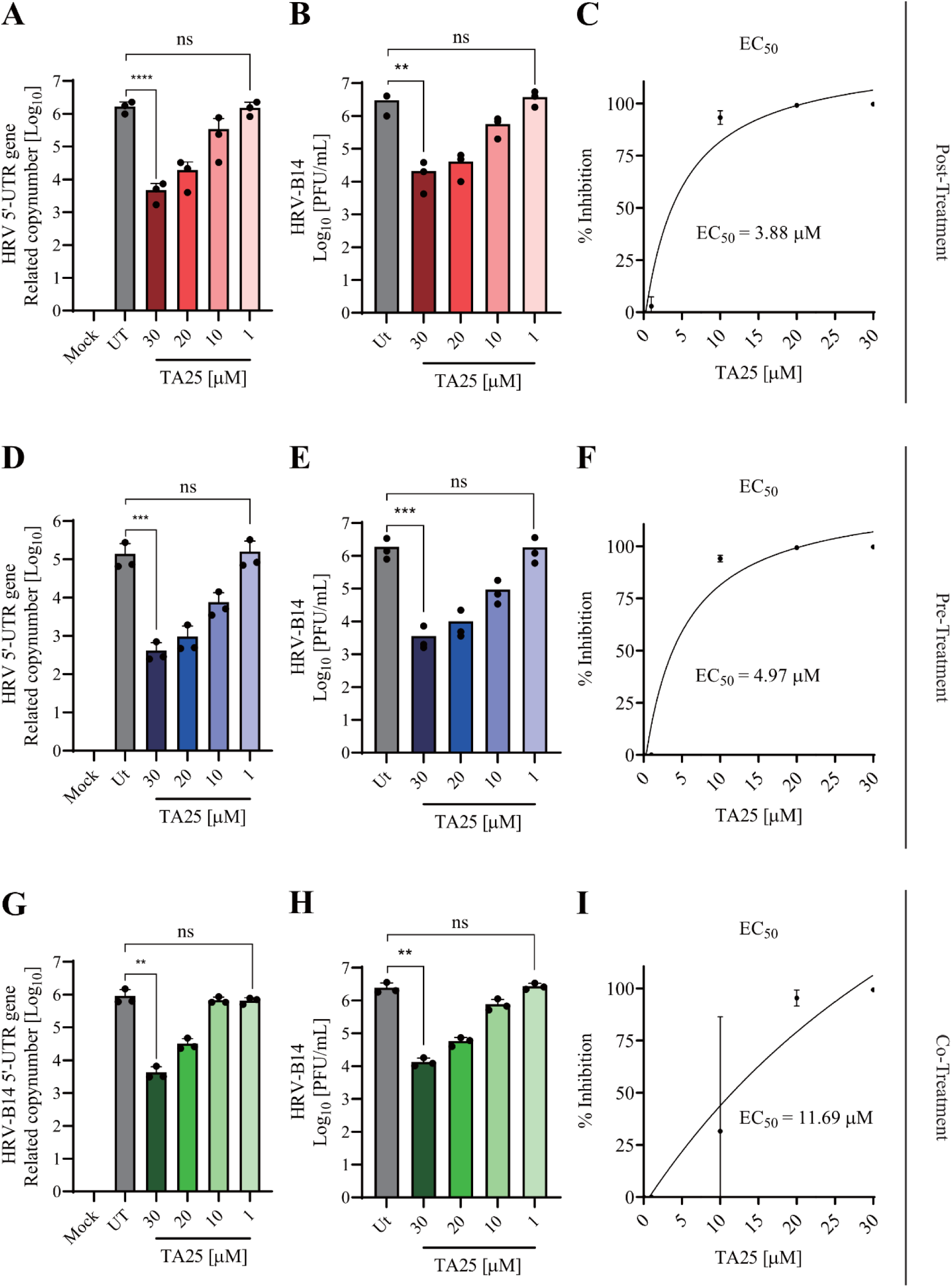
Antiviral activity of post-, pre-, and co-treatment of TA25 against HRV-B14. Antiviral efficacy of TA25 against HRV-B14 infection in HeLa cells after post-treatment, pre-treatment, and co-treatment. Cells were infected with HRV-B14 at an MOI of 0.1 and treated with 30, 20, 10, and 1 μM of TA25. Supernatants for plaque assays and infected cells for RT-qPCR were harvested at 48 h post-infection (hpi). (A), (D), and (G) Expression levels analyses of HRV-B14 5’-UTR genes in HeLa cells using RT-qPCR under post-, pre-, and co-treatment conditions, respectively. (B), (E), and (H) Virus titres of HRV-B14 determined by plaque assays post-, pre-, and co-treatment, respectively. (C), (F), and (I) EC_50_ values of TA25 against HRV-B14 after post-, pre-, and co-treatment, respectively. The data presented are representative of three independent experiments performed in triplicate. *p < 0.05, **p < 0.01, ***p < 0.001, and **** p < 0.00001, t-tests (Ut = untreated), (ns = not-significant).

### 2.2. TA25 reduces the expression of innate immune and pro-inflammatory genes induced by HRV-B14

To investigate the effect of TA25 on innate immune and pro-inflammatory response, total RNA-seq and RT-qPCR analyses were performed on mock, HRV-B14-infected, and 30 μM of TA25 treated HeLa cells. RNA-seq revealed an induction of interferon (IFN)-related and pro-inflammatory genes following HRV-B14 infection. In contrast, the expression of IFN-related and pro-inflammatory genes was reduced in TA25-treated cells (Figures 2A and B). RT-qPCR validation confirmed the decreased expression of RIG-I, IFN-β, ISG15, ISG54, and ISG56 in TA25-treated cells compared with infected controls (Figure 2C). In addition, pro-inflammatory markers including TNFAIP2, TNFAIP6, CXCL10, CXCL11, and STAT1 were significantly downregulated in the TA25-treated cells (Figure 2D). These findings suggest that TA25 suppresses the host innate immune and pro-inflammatory genes responses triggered by HRV-B14 infection.

**Figure 2.**
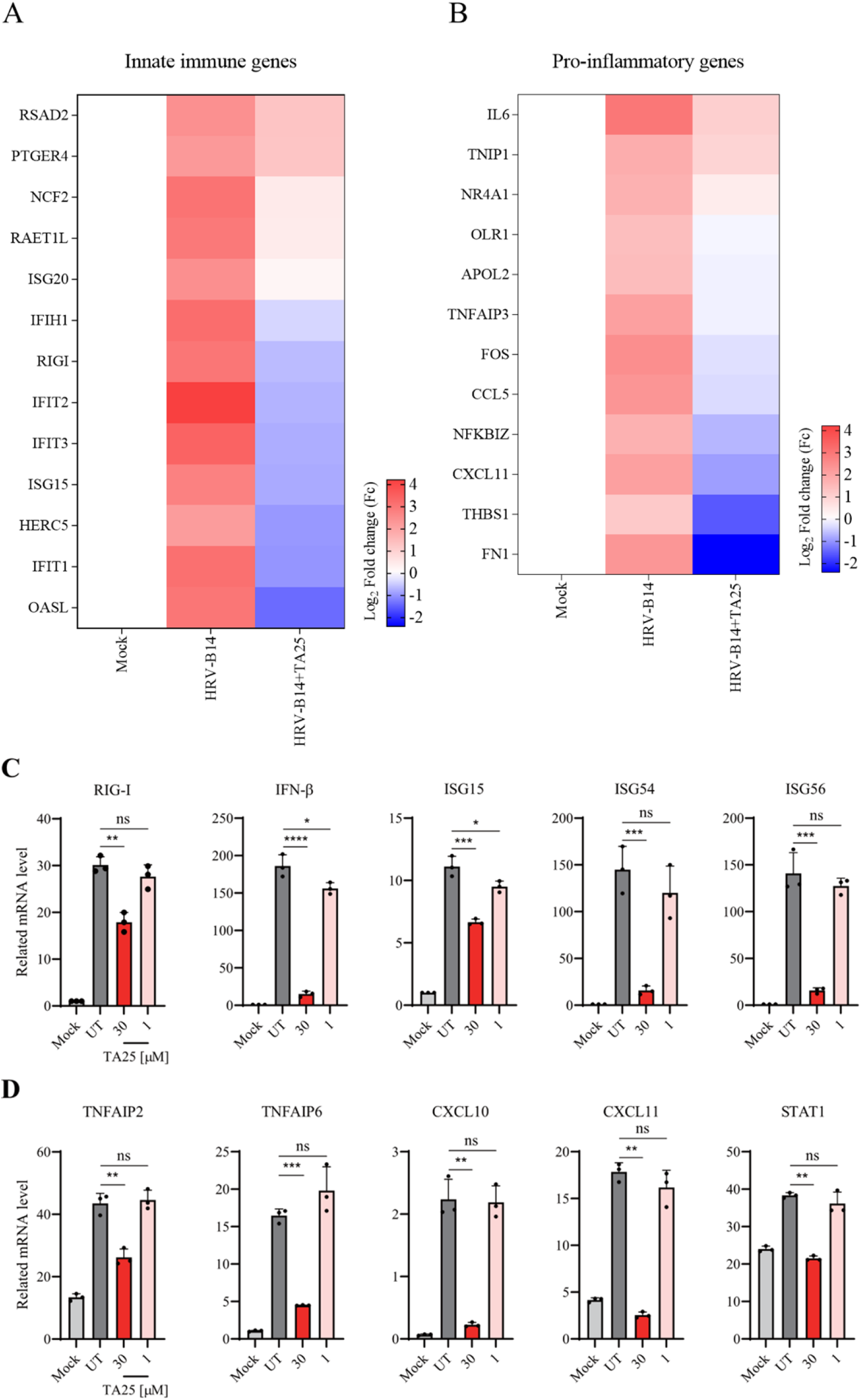
Down-regulation of innate immune and pro-inflammatory genes after TA25 treatment of HRV-B14 infected HeLa cells. To analyze TA25’s effect on innate immune and pro-inflammatory response genes produced by HRV-B14 infection, the changes in expression levels were compared between HRV-B14-infected cells and cells treated with 30 μM TA25 2 h after HRV-B14 infection and cells were incubated for 48 h. Total RNA-seq was performed using NovaSeq 6000, and DEGs were confirmed by RT-qPCR. (A) Heatmap of innate immune response genes in HRV-B14-infected and TA25-treated cells. (B) Heatmap of pro-inflammatory response genes in HRV-B14-infected and TA25-treated cells. (C) RT-qPCR analysis of IFN-related pathway genes and (D) pro-inflammatory genes in HRV-B14-infected and TA25-treated cells. The data presented are representative of three independent experiments performed in triplicate. *p < 0.05, **p < 0.01, ***p < 0.001, and **** p < 0.00001, t-tests (Ut = untreated), (ns = not-significant).

### 2.3. TA25 inhibits the synthesis of genomic and anti-genomic HRV RNA

To evaluate the inhibitory effects of TA25 on the viral protein translation and/or viral RNA replication, HRV strand-specific RT-qPCR was performed on 30 μM of TA25-treated HeLa cells at 6, 12, 24, and 48 hpi. At 6 hpi, genomic RNA was detected. At 6 hpi, replication had not yet occurred, but early translation to synthesize proteins for replication appeared to occur (Figure 3A and B). At 12 hpi, anti-genomic RNA began to be amplified only in virus-infected cells and decreased in TA25-treated cells. RNA synthesis did not increase significantly indicated by a relative expression level of 1. Similarly, the relative expression levels of 12 and 24 hpi were not increased significantly with 6.4 and 6.9, respectively. The anti-genomic RNA was not synthesized using the genomic RNA as a template, and the synthesis of the genomic RNA using anti-genomic RNA was decreased in TA25-treated cells. (Figure 3B). This suggests that TA25 inhibited viral genomic amplification.

**Figure 3.**
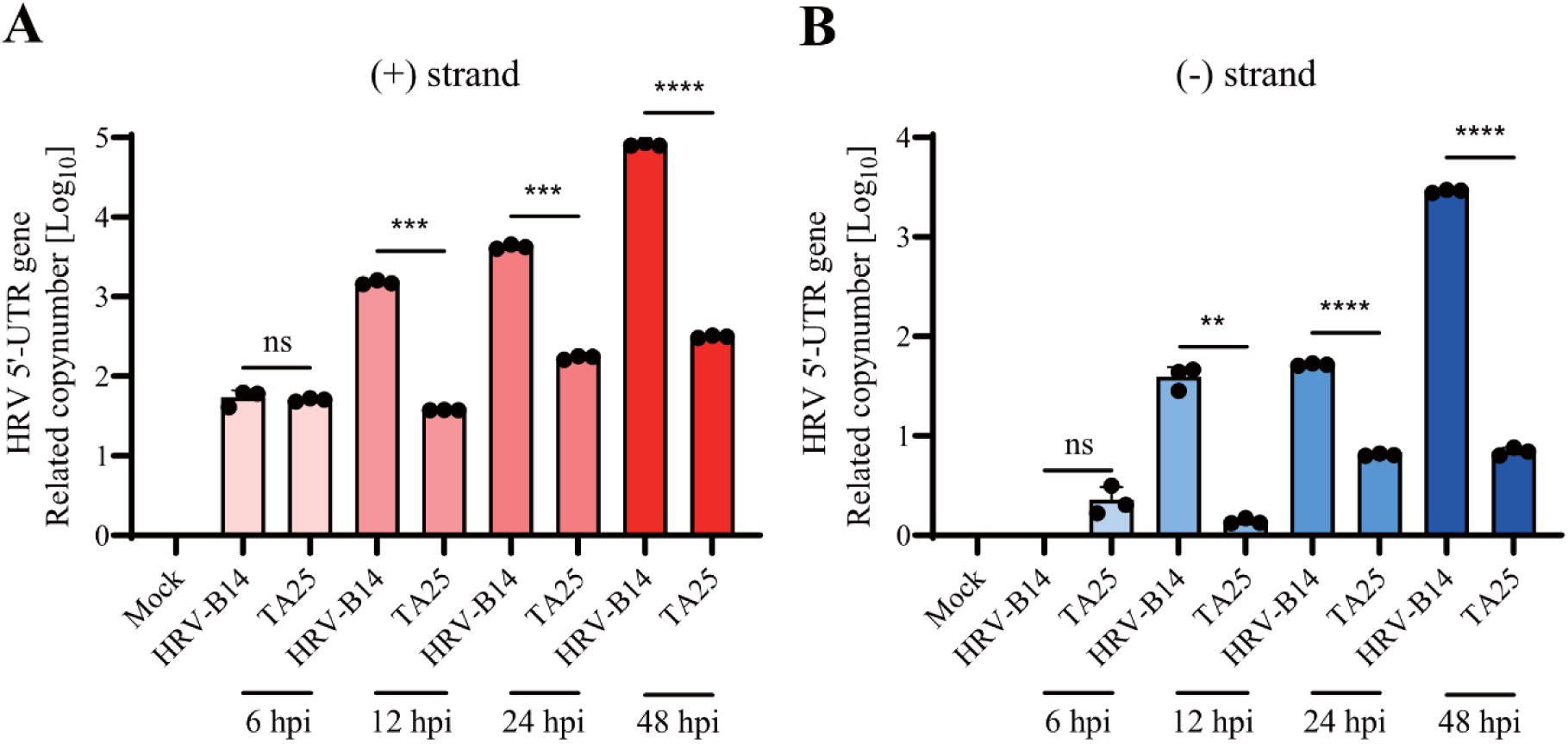
Replication of human rhinovirus is inhibited in TA25-treated cells. HRV-B14 strand-specific RT-qPCR was performed at different hpi to determine when the viral genome decreased in size during the viral life cycle. HeLa cells were treated with 30 μM of TA25 for 2 h after HRV-B14 infection at an MOI of 0.1. (A) genomic RNA and (B) anti-genomic RNA expression levels of HRV-B14 at 6, 12, 24, and 48 hpi were measured by RT-qPCR. The data presented are representative of three independent experiments performed in triplicate. *p < 0.05, **p < 0.01, ***p < 0.001, and **** p < 0.00001, t-tests (Ut = untreated), (ns = not-significant).

### 2.4. AI-driven structure-based docking analyses exhibit the interaction of TA25 with 2B protein

To identify the molecular targets of TA25, AI-based docking tests between TA25 and HRV-B14 were conducted using the Pharmaco-Net platform developed by CALICI. As shown in Figure 4A, TA25 was predicted to bind to the active site of the 2B protein. The predicted binding energy and binding affinity of TA25 to the 2B protein were estimated to be –4.2 kcal/mol and 0.0074 μM, respectively (Figure 4B). To further identify the potential targets, additional docking analyses revealed the interaction of TA25 with RdRp and 3C protease proteins. The predicted binding energies for TA25 with RdRp and 3C protease were –4.74 kcal/mol and –5.23 kcal/mol, respectively, with corresponding predicted binding affinities of 0.974 μM and 1.56 μM (Table S1). The binding affinity of TA25 for the 2B protein was significantly stronger than that of the RdRp and 3C proteases, suggesting that TA25 targets 2B protein to inhibit HRV replication.

**Figure 4.**
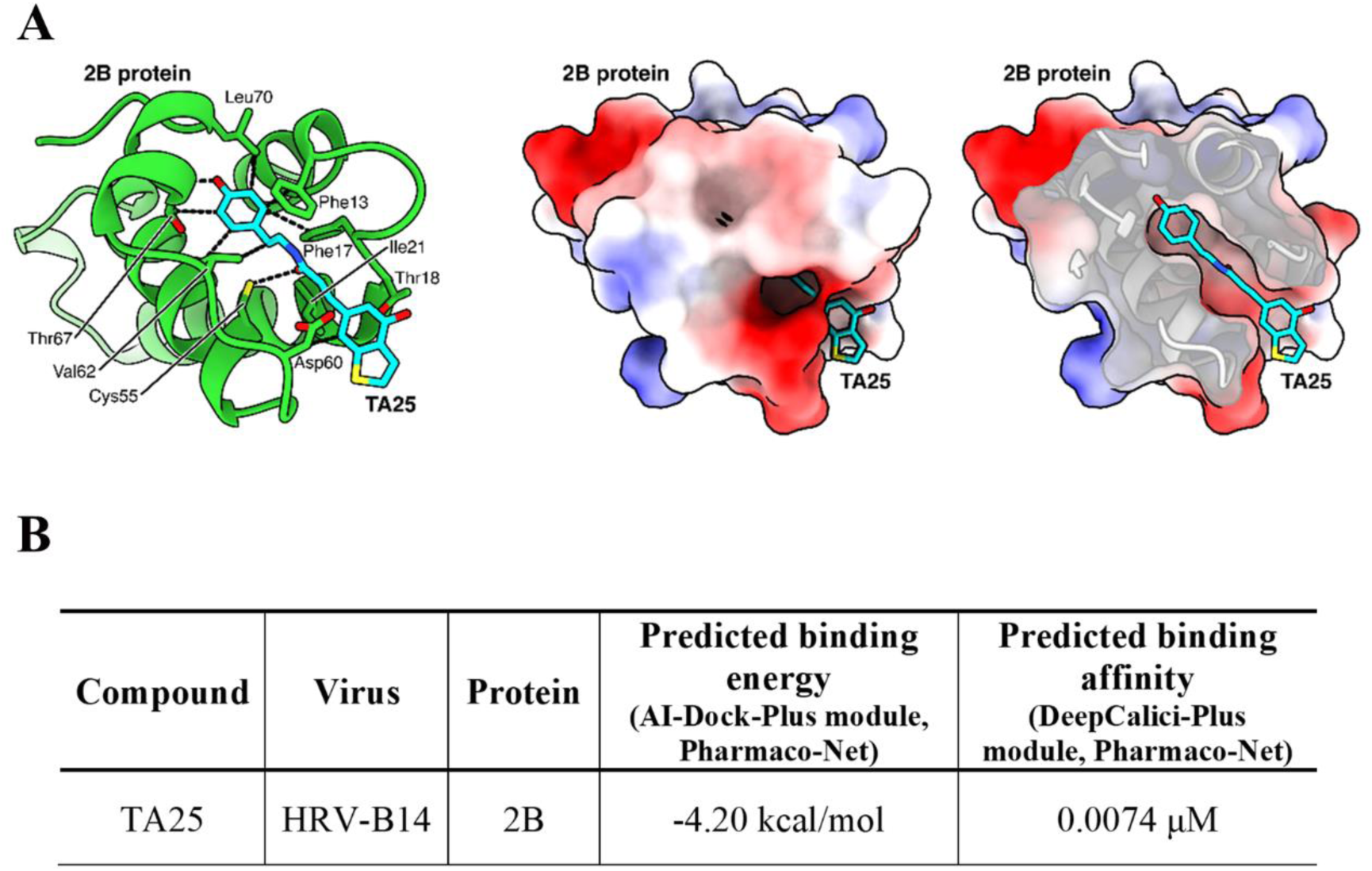
AI-driven structure-based docking analysis of TA25 and 2B protein of HRV-B14. To identify the interactions between TA25 and HRV protein, an AI-driven structure-based analysis was performed. Key residues of the 2B protein predicted to interact with TA25 were indicated (left). Electrostatic surface views were shown (middle and right). (A) The predicted pose of TA25 with the HRV 2B protein was identified computationally. (B) The predicted binding energy and affinity for TA25 binding to 2 B protein were calculated using the AI-Dock-Plus module and DeepCalici-Plus modules, respectively, in Pharmaco-Net.

### 2.5. TA25 confers broad-spectrum antiviral activities against *Picornaviridae*, *Flaviviridae*, *Poxviridae*, and *Orthomyxoviridae*

To test the broad-spectrum antiviral activity of TA25, antiviral screening was performed against multiple viruses including HRV-1A (*Picornaviridae*), ZIKV (*Flaviviridae*), DENV-2 (*Flaviviridae*), VACV (*Poxviridae*), IBV Victoria (*Orthomyxoviridae*), and RSV-A (*Pneumoviridae*). The VP1 genes of HRV-1A were reduced with approximately 100-fold at the RNA expression levels (Figure 5A). After 48 hpi, the relative expression of ZIKV NS1 genes was reduced approximately 100-fold with 30 μM of TA25 (Figure 5B). The TA25 treatments showed an approximately 50-fold reduction in DENV-2 E genes expression (Figure 5C). In the case of VACV, E9 genes expression levels were decreased by approximately 50-fold (Figure 5D). The expression of IBV Victoria HA genes was decreased approximately 6-fold when MDCK cells were treated with 30 μM of TA25 (Figure 5E). In contrast, TA25 showed no antiviral activity against RSV-A in HEp-2 cells (Figure 5F). The EC_50_ value for HRV-1A, ZIKV, DENV-2, IBV Victoria and VACV were 9.7 μM, 1 μM, 5.6 μM, 15.3 μM and 1.7 μM, respectively (Table S2). These results suggest that TA25 exerts broad-spectrum antiviral activity across both RNA and DNA viruses.

**Figure 5.**
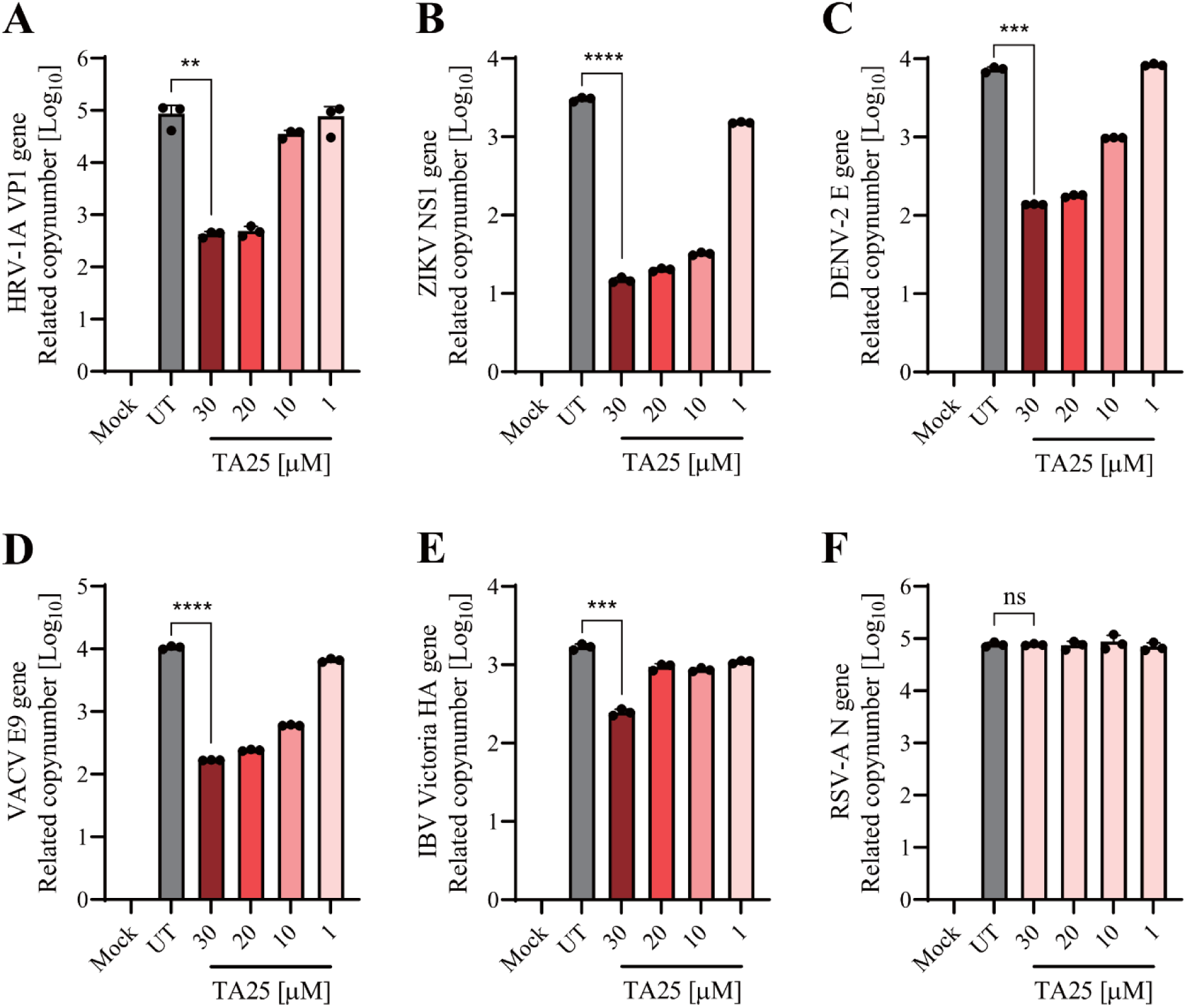
Antiviral efficacy of TA25 against multiple zoonotic viruses. To evaluate the broad-spectrum antiviral activity of TA25, its antiviral activity was tested against ZIKV, DENV, VACV, IBV Victoria and RSV. 2 h after infection, the cells were treated with 30, 20, 10, and 1 μM of TA25 and total RNA was collected at 48 hpi. The expression of (A) HRV-1A VP1, (B) ZIKV NS1, (C) DENV E, (D) VACV E9, (E) IBV Victoria HA, and (F) RSV N genes was analyzed in HeLa cells. HeLa cells were infected with HRV-1A; Vero E6 cells were infected with ZIKV, DENV, or VACV; MDCK cells were infected with IBV Victoria; and Hep-2 cells were infected with RSV. The data presented are representative of three independent experiments performed in triplicate. *p < 0.05, **p < 0.01, ***p < 0.001, and **** p < 0.00001, t-tests (Ut = untreated), (ns = not-significant).

## 3. Discussion

HRV infects both the upper and lower respiratory tracts (18). HRV infection aggravates pre-existing lung conditions including asthma, pneumonia, and COPD (19). Currently, no antiviral drugs or vaccines are approved for the prevention of HRV infection (20). Several candidate antiviral compounds, such as rupintrivir (AG-7088), have been identified against HRV (21); however, their efficacy in treating natural infections has been limited (22). To evaluate the potential of TA25 as an anti-HRV agent, its effect was tested in HRV-B14 infected cells under three conditions: post-treatment, pre-treatment, and co-treatment (Figure 1). In all three conditions, TA25 reduced viral replication by approximately 100-fold with minimal cytotoxicity (Figure S1). As shown in Figure 2, TA25 significantly down-regulated the expression of innate immune- and proinflammation-related genes in HRV-B14-infected cells. These findings suggest that TA25 may serve as a potential antiviral and anti-inflammatory agent.

Numerous replication inhibitors have been commercialized for respiratory viruses (23). Remdesivir, the first FDA-approved antiviral for SARS-CoV-2, exhibits broad-spectrum antiviral activity against *Betacoronaviruses* including SARS-CoV-2, SARS-CoV and MERS-CoV, Enterovirus, Ebolavirus and RSV (24). The active form of remdesivir functions as a nucleoside analog, inhibiting the viral RdRp (25). Remdesivir demonstrated the antiviral activity against SARS-CoV-2 infection in primary human airway epithelial cells with an EC_50_ of 9.9 nM at 48 hpi (26). A previous study showed that TA25 inhibited influenza A virus (IAV) replication in A549 cells at 4 and 8 hpi (27). The synthesis of NS1 positive-sense RNA was significantly reduced at 4 and 8 hpi, suggesting that the TA25 might inhibit the transcription or replication of IAV. In this study, strand-specific RT-qPCR of HRV genomes showed that at 6 hpi, the positive-sense RNA, representing the genomic RNA, had not yet been replicated into the anti-genomic RNA (Figure 3). By 12 hpi, the anti-genomic RNA began to be synthesized; however, its amplification was inhibited over time. As a result, the genomic RNA could not be effectively replicated using the anti-genomic RNA as a template. This finding suggests that TA25 may bind to viral proteins associated with the replication, leading to the inhibition of anti-genomic RNA synthesis. Using the AI-based drug discovery platform, Pharmaco-Net, we found most appropriate binding affinity and binding energy of TA25 with HRV 2B protein compared with RdRp and 3C protease. The previous study demonstrated that the 2B protein facilitated HRV replication by interacting with host proteins and phospholipids to form replication organelles (ROs) (28). The 2B protein led to the release of the immune adaptor protein STING from the endoplasmic reticulum and targeted it to ROs enriched with the phosphatidylinositol 4-phosphate (PI4P). The replication complex, consisting of viral proteins 2B, 2C, and 3A, played a key role in the synthesis of nascent RNA with STING and PI4P in these ROs. However, the precise mechanism of action of TA25 remains to be further investigated.

Broad-spectrum antiviral agents are compounds capable of inhibiting infections caused by multiple viruses, either within the same or across different families (29). Virus- or host-direct antivirals are limited to emerging and re-emerging viruses against which no vaccines or antiviral therapies are approved (30). Broad-spectrum therapeutics could address this issue by being administered before a viral threat has been accurately diagnosed, thereby controlling the spread of virus. Emerging and re-emerging viruses, including DENV, West Nile Virus (WNV), ZIKV, SARS-CoV-2, and influenza virus, pose significant public health concerns (31). Currently, international travel, migration, and the global commerce facilitate the spread of emerging viruses, increasing the risk of epidemic and pandemic outbreaks (32). There is a significant unmet need for the development of broad-spectrum antiviral therapeutics for the emerging and re-emerging viral infections. Moreover, TA25 was effective in limiting the replication of ZIKV, DENV, and VACV, leading to the hypothesis that this antiviral agent may act by interfering with viral genome synthesis (Figure 5). These findings suggest the potential of TA25 as a broad-spectrum antiviral capable of inhibiting the replication and spread of multiple emerging viruses.

## 4. Conclusion

This study demonstrates the antiviral activity of TA25 against HRVs and multiple zoonotic viruses *in vitro* and *in silico*. The antiviral efficacy of TA25 appears to be mediated by inhibiting HRV genome replication. Thus, these findings provide insight into the potential of TA25 for the development of novel antiviral therapeutics to mitigate the replication of emerging RNA and DNA viruses.

## 5. Materials and Methods

### 5.1. Ethics

The study of antiviral reagents against HRV-B14, HRV-1A, ZIKV, DENV-2, VACV, IBV Victoria, and RSV-A was performed at biosafety level-2, following the guidelines and protocols of the Hallym University Institutional Biosafety Committee.

### 5.2. Cell culture

HeLa, Vero E6, MDCK and HEp-2 cells were purchased from American Type Culture Collection (ATCC), were cultured in Dulbecco’s Modified Eagle’s Medium (DMEM; Cat No. 11965092, Gibco), supplemented with 10% Fetal Bovine Serum (FBS; Cat No. 16000044, Gibco), 1% HEPES (1 M; Cat No. 15630080, Gibco) and 1% Antibiotic-Antimycotic (Anti-Anti; 100 X; Cat No. 15240062, Gibco). Cells were incubated at 37℃ with 5% CO_2_.

### 5.3. Viruses

HRV-B14 (ATCC VR-284) and HRV-1A (ATCC VR-1559) were acquired from the ATCC. ZIKV (NCCP No. 43280), DENV-2 (NCCP No. 43248), VACV (NCCP No. 43281) and IBV Victoria (NCCP No. 43402) were acquired from the National Culture Collection for Pathogens (Osong, Republic of Korea) and RSV-A (KBPV-VR-41) was acquired from the Korea Bank for Pathogenic Viruses for use in this study.

### 5.4. *In vitro* infection and TA25 treatment

HeLa, Vero E6, MDCK and HEp-2 cells were seeded in 6-well plates (Cat No. 353046, Falcon) to reach 80% confluence and incubated overnight. Next day, the cells were washed twice with pre-warmed phosphate-buffered saline (PBS) (Cat No. 70011069, Gibco). Three experimental conditions were established: post-, pre-, and co-TA25 treatment. For post-treatment, HeLa cells were infected with HRV-B14 and HRV-1A at a multiplicity of infection (MOI) of 0.1 at 34℃ with 5% CO_2_. Vero E6 cells were infected with ZIKV, DENV-2 and VACV at a MOI of 0.01, MDCK cells were infected with IBV Victoria at a MOI of 0.01 and HEp-2 cells were infected with RSV-A at a MOI of 0.1 at 37℃ with 5% CO_2_. The plates were shaken every 15 min for 2 h to efficiently distribute the viruses. After 2 h of infection, the cells were treated with TA25 (339.4 g/mol) at different concentrations. For pre-treatment, HeLa cells were treated with TA25 at different concentrations for 2 h and infected with HRV-B14 at an MOI of 0.1. For co-treatment, HeLa cells were infected with HRV-B14 at an MOI of 0.1 and treated with TA25 at different concentrations at the same time. Supernatants and infected cells were harvested at 48 h post-infection (48 hpi) for analysis.

### 5.5. RNA extraction and reverse transcription-polymerase chain reaction (RT-PCR)

Total RNA from the infected cells was collected using 1 mL of TRIzol reagent (Cat No. 15596026, Invitrogen) according to the manufacturer’s protocol. RNA concentration and quality were assessed using a NanoDrop Spectrophotometer (Cat No. ND-2000, Thermo Scientific, NanoDrop 2000/2000c). 1 μg of RNA was reverse-transcribed into cDNA using a High-Capacity RNA-to-cDNA kit (Cat No. 4387406, Applied Biosystems) with a SimpliAmp Thermal Cycler (Cat No. A24811, Applied Biosystems). cDNA synthesis for strand-specific RT-PCR was performed using 1 μM of genomic or anti-genomic specific RT-PCR primers that are listed in Table S3 using the same PCR program.

### 5.6. Real-Time quantitative polymerase chain reaction (RT-qPCR)

To quantify viral and host gene expression levels, RT-qPCR was performed using Power SYBR Green PCR Master Mix (Cat No. 4367659, Applied Biosystems) on a QuantStudio3 real-time PCR instrument (Cat No. A28567, Applied Biosystems). RT-qPCR was performed as follows: initial denaturation at 50℃ for 2 min and 95℃ for 10 min; 40 PCR cycles at 95℃ for 15 s and 60℃ for 1 min (65℃ for SARS-CoV-2 and 47℃ for strand-specific RT-qPCR); elongation at 95℃ for 15 s and 65℃ for 1 s. Primer sequences for strand-specific RT-qPCR, viral and human-related genes are listed in Tables S3, 4 and 5, respectively.

### 5.7. Plaque assay

HeLa cells (1×10^6^ cells / well) were seeded in a 6 well plate and incubated to achieve 100% confluency. The cells were washed with pre-warmed PBS and infected with 10-fold diluted HRV-B14 supernatant in serum-free DMEM at 34°C with 5% CO_2_ for 4 h. The plates were gently shaken every 15 minutes. The cells were covered with 0.6% oxoid agarose containing overlay EMEM media containing 5% 1 M HEPES, 2% FBS, 2% 100 × non-essential amino acids (NEAA) (Cat No. 11140050, Gibco), 2% 200 mM L-glutamine (Cat No. 25030081, Gibco), and 0.2% gentamycin (50 mg/mL) (Cat No. 15750078, Gibco) for four days. The cells were fixed with 3.7% formaldehyde (Cat No. F1119Z21YR; Biosesang) and stained with 0.1% crystal violet in 20% methanol.

### 5.8. Total RNA-sequencing (Total RNA-seq) analysis

RNA quality was assessed using a TapeStation4000 System (Cat No. G2991BA, Agilent) for library preparation of total RNA sequencing data. rRNA was depleted using the RiboCrop rRNA depletion kit (Cat No. 144, LEXOGEN), the cDNA library was prepared using the CORALL RNA-seq V2 Library Prep Kit (Cat No. 0001-0096, LEXOGEN), sequencing was performed by NovaSeq 6000 analysis, and statistical processing was performed after mapping with STAR to the reference genome, and mapped reads were calculated using salmon. Transcript abundance was normalized to count per million (CPM) mapped reads. The TMM method estimates the scale factors between samples for differentially expressed gene (DEG) analysis. DEG analysis compares the differences in gene expression and regulation patterns between sample groups after transcriptome sequencing of two or more samples.

### 5.9. Cell counting kit-8 assay

The cytotoxicity of TA25 was determined using a Cell Counting Kit-8 (CCK-8) assay (Dojindo Molecular Technologies, Kumamoto, Japan) following the manufacturer’s protocol. Briefly, cells were plated in 96-well plates (10^4^ cells / 100 μL / well) and incubated for 24 h before TA25 treatment. The next day, cells were treated with different concentrations of TA25 at 34°C and 5 % CO_2_ for 48 h. At 45 h post treatment, 10 μL of CCK-8 reagent was added to the wells. After 3 h, the optical density (OD) was measured using a microplate reader at 450 nm.

### 5.10. AI-driven structure-based docking analysis

Structure-based docking simulations between compound TA25 and HRV proteins were performed using the Pharmaco-Net platform (https://pharmaco-net.org), which integrates advanced AI-powered tools, including PocketFinder, AI-Dock-Plus, DeepCalici-Plus, and InteractionViewer modules developed by CALICI (https://calici.co). The three-dimensional (3D) structure of the HRV 2B protein was predicted using SWISS-MODEL based on its amino acid sequence (NP_740520.1). The experimentally resolved 3D structures of RdRp (PDB ID: 1XR5) and 3C protease (PDB ID: 5FX6) were retrieved from the Protein Data Bank. Following the upload of the 3D structures of 2B, RdRp, and 3C proteases, potential ligand-binding sites were identified using the PocketFinder module. Subsequently, the 3D structure of TA25 was uploaded, and its predicted binding energy and binding affinity to each viral protein were computed using AI-Dock-Plus and DeepCalici-Plus modules, respectively. Finally, the spatial binding position of TA25 within the predicted active sites of each target protein was visualized using the InteractionViewer module of Pharmaco-Net. Structural figures were generated using UCSF ChimeraX (33).

### 5.11. Statistical analysis

Statistical analyses were performed using GraphPad Prism (Version 8.0.2; GraphPad Software, Inc., La Jolla, CA, USA). Values are presented in bar graphs as the mean ± SD from at least three independent experiments; *p < 0.05, **p < 0.01, ***p < 0.001, and **** p < 0.0001 were considered statistically significant.

## Abbreviations

COPD: chronic obstructive pulmonary disease
DENV: dengue virus
HRV: Human rhinovirus
IBV: influenza B virus
RSV: respiratory syncytial virus
TA25: Tabamide A 25
VACV: vaccinia virus
ZIKV: zika virus

## Declaration of competing interest

The authors declare no competing financial interests.

## Acknowledgements

We thank Ms. Nayeon Jang, Biomedical Research Institute, Hallym University, School of Medicine, for her supporting this study.

## Author contributions: CRediT

**Bohyeon Kim:** Conceptualization, Data curation, Methodology, Writing – original draft. **Altanzul Bat Ulzii:** Formal analysis, Data curation. **Taehun Lim:** Validation, Data curation. **Jin Woo Kim:** Data curation, Resources. **Wonkyun Ronny Im:** Data curation, Resources. **Shivani Rajoriya:** Data curation, Methodology. **Augustine Natasha:** Data curation, Methodology. **Divya Misra:** Data curation. **Hennisa Hennisa:** Data curation. **Na Young Kwag:** Data curation. **Daewoon Yoon:** Data curation. **Gyunghee Jo:** Data curation, Writing – review & editing. **Young Bin Park:** Resources, Writing – review & editing. **Jinkyu Park:** Conceptualization, Validation. **Jeong Tae Lee:** Resources, Conceptualization. **Sung Ho Jeon:** Resources, Conceptualization. **Won-Keun Kim:** Conceptualization, Funding acquisition, Supervision, Writing – review & editing.

## Funding

This research was supported by the Korea Institute of Marine Science and Technology Promotion (KIMST), funded by the Ministry of Oceans and Fisheries, Korea (RS-2021-KS211475) and the Ministry of Education and National Research Foundation of Korea, funded by the Glocal University (GLOCAL-202504590001).

## Supplementary Figure

**Supplementary Figure S1.**
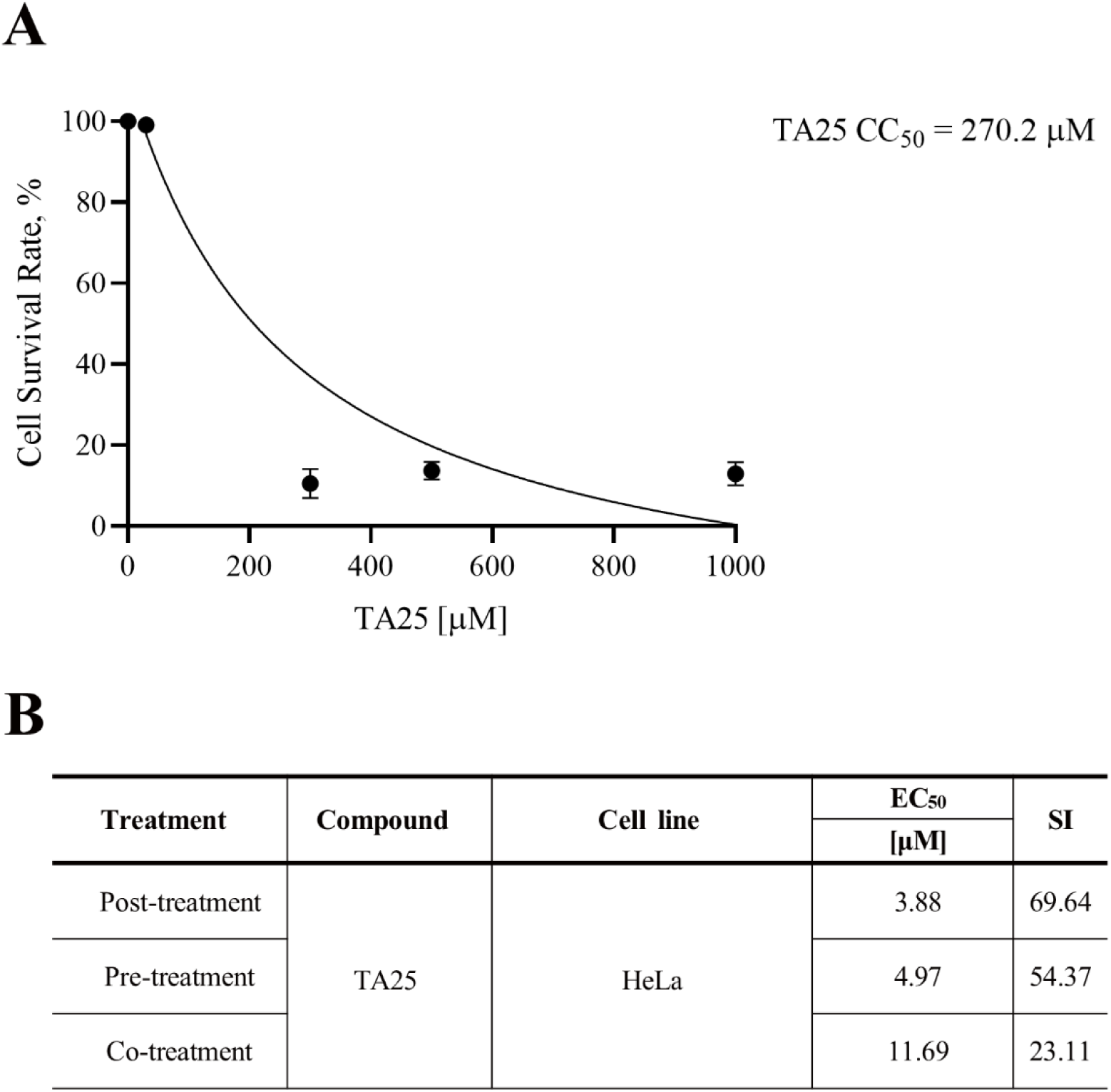
Evaluation of TA25 cytotoxicity in HeLa cells at a maximum of 1,000 μM. The cytotoxicity of TA25 was tested in HeLa cells incubated for 24 h with concentrations of 30, 100, 300, 500, and 1,000 μM TA25. NADH was detected using an absorbance wavelength of 450 nm. (A) Half-maximal cytotoxic concentration for HeLa cells. (B) Selectivity index (SI), the ratio of to CC_50_/EC_50_. The data presented are representative of three independent experiments performed in triplicates.

## Supplementary Tables

**Table S1.** AI-driven structure-based analyses of TA25 with HRV-B14 RdRp and 3C protease.

| Compound | Virus | Protein | Predicted binding energy<br>(AI-Dock-Plus module,<br>Pharmaco-Net) | Predicted binding affinity<br>(DeepCalici-Plus<br>module, Pharmaco-Net) |
| --- | --- | --- | --- | --- |
| TA25 | HRV-B14 | RdRp | -4.74 kcal/mol | 0.974 μM |
| | | 3C<br>protease | -5.23 kcal/mol | 1.56 μM |

**Table S2.** Mean of effective concentration 50 (EC_50_) of TA25 against different strains of human rhinovirus and multiple viruses.

| Treatment | Type | Family | Viruses | EC<sub>50</sub> |
| --- | --- | --- | --- | --- |
|  |  |  |  | [μM] |
| TA25 | RNA | *Picornaviridae* | HRV-1A | 9.717 |
|  |  | *Flaviviridae* | ZIKV | 1.077 |
|  |  | *Flaviviridae* | DENV-2 | 5.665 |
|  |  | *Orthomyxoviridae* | IBV Victoria | 15.32 |
|  | DNA | *Poxviridae* | VACV | 1.732 |

**Table S3.**
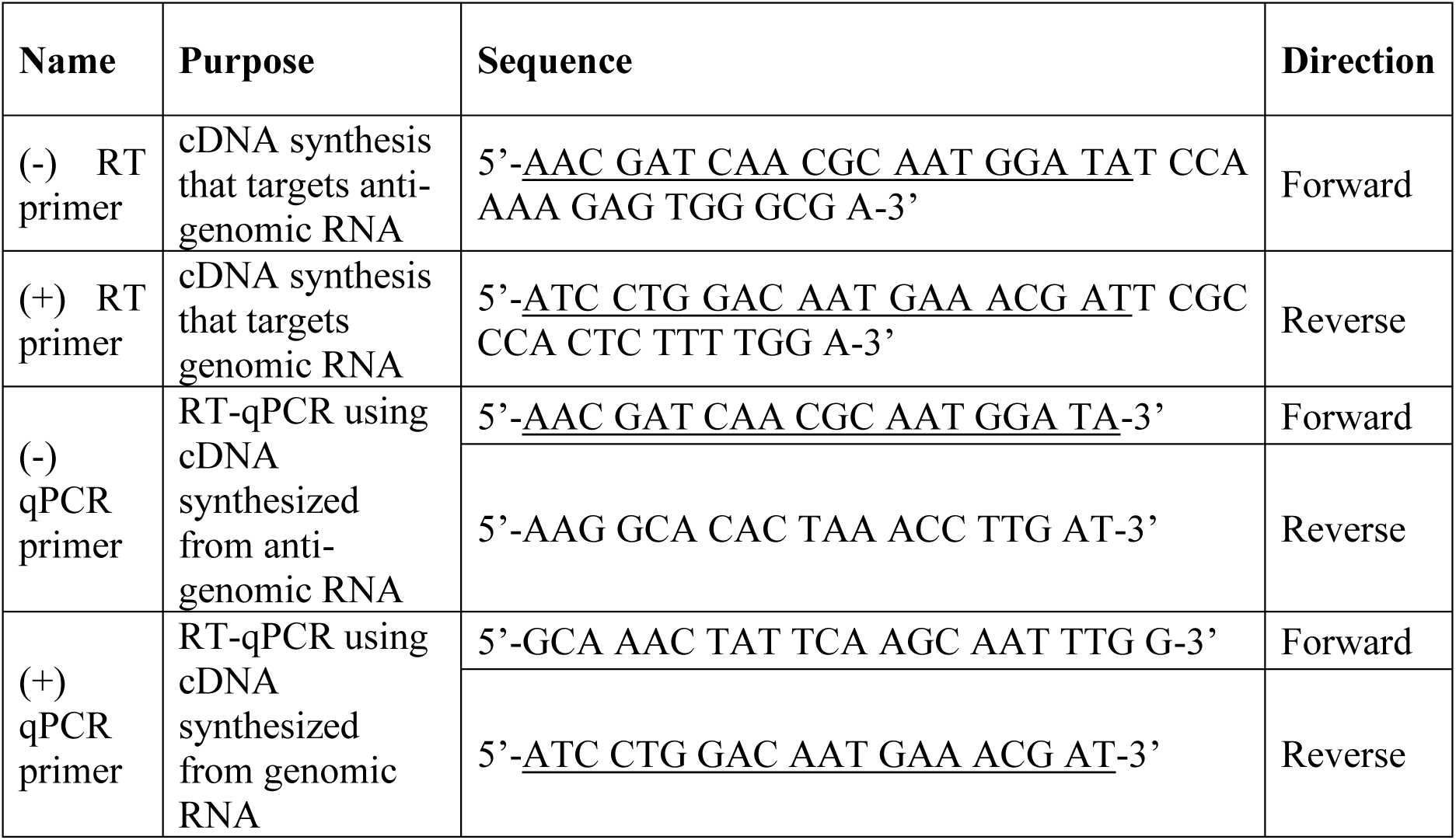
Primer sequences of strand-specific RT-PCR and RT-qPCR.

**Table S4.**
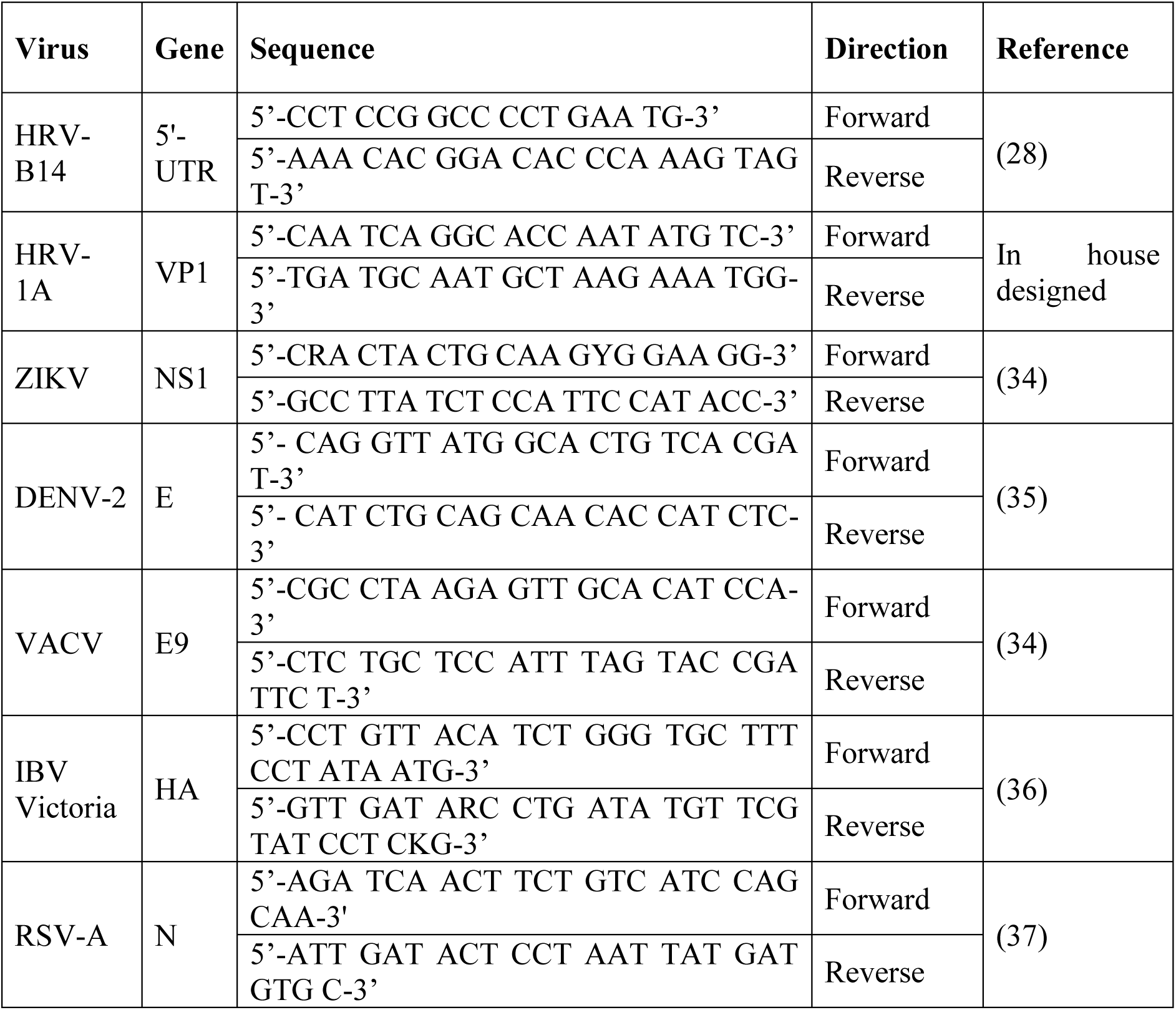
Primer sequences of viral genes for RT-qPCR.

**Table S5.**
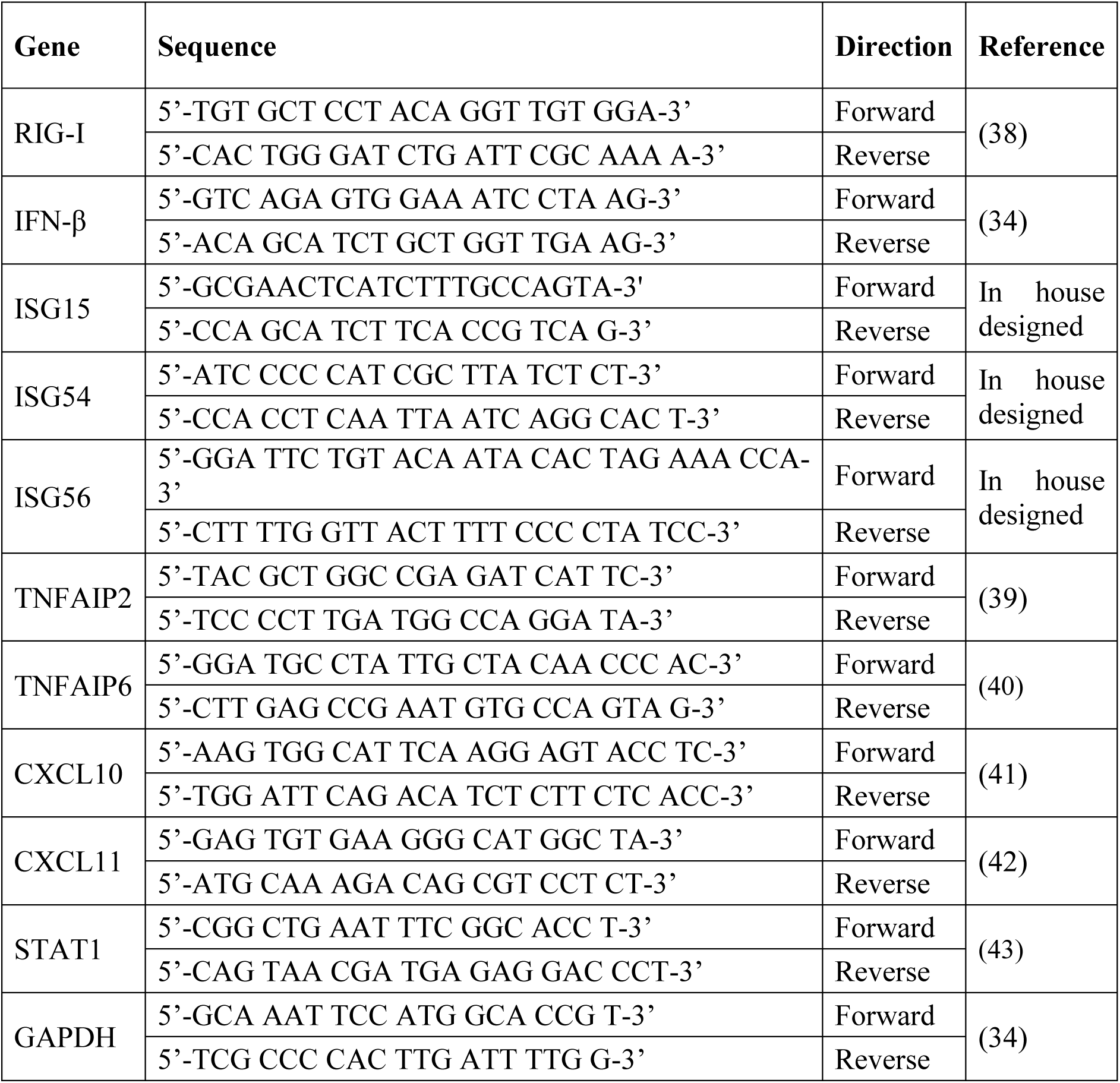
Primer sequences of human IFN-related and inflammatory genes for RT-qPCR.

